# Automatic identification of scientific publications describing digital reconstructions of neural morphology

**DOI:** 10.1101/2023.02.14.527522

**Authors:** Patricia Maraver, Carolina Tecuatl, Giorgio A. Ascoli

## Abstract

**Motivation:** The increasing number of peer-reviewed publications constitutes a challenge for biocuration. For example, NeuroMorpho.Org, a sharing platform for digital reconstructions of neural morphology, must evaluate more than 6000 potentially relevant articles per year to identify data of interest. Here, we describe a tool that uses natural language processing and deep learning to assess the likelihood of a publication to be relevant for the project.

**Results:** The tool automatically identifies articles describing digitally reconstructed neural morphologies with high accuracy. Its processing rate of 900 publications per hour is not only amply sufficient to autonomously track new research, but also allowed the successful evaluation of older publications backlogged due to limited human resources. The number of bio-entities found since launching the tool almost doubled while greatly reducing manual labor. The classification tool is open source, configurable, and simple to use, making it extensible to other biocuration projects.

**Availability:** https://github.com/Joindbre/TextRelevancy

**Supplementary information:** Supplementary information, tool installation, and API usage are available at https://docs.joindbre.com

## 1 Introduction

Biocuration is the conversion of peer reviewed biomedical information into accessible data understandable by humans and machines (Howe *et al*., 2008). The exponential growth of the scientific literature makes text mining an indispensable element of the biocuration workflow (Hirschman *et al*., 2012). The process involves multiple operations: finding the relevant publications (triage), identifying the bio-entity data within the text, extracting and normalizing the selected data, storing the entries into a database, and validating the correctness of the re-structured information.

Triage is one of the most labor-intensive tasks: biocurators use search engines to query potentially relevant publications using keywords; they must then inspect the resultant collection of articles to identify those containing information that needs to be extracted.

Multiple machine learning approaches have been applied to reduce the effort expended in triage. When employing shallow methods like Naïve Bayes and Random Forest on input consisting of the article title, abstract, and figure captions, use of balanced datasets generally outperforms imbalanced datasets (Jiang *et al*., 2017), and the same holds true for Support Vector Machines algorithm (Jiang *et al*. 2020, 2019). A more comprehensive study reviews performance varying the imbalanced proportions, gradually under-sampling the majority class by a factor of 5% from 90-10% to 50-50% (Almeida *et al*., 2014); although recall improved with balance, it was close to random for all the tests performed. In addition, deep methods like Convolutional Neural Networks (CNN) (LeCun *et al*., 1989) have been combined with word embeddings (Bengio *et al*., 2003) to capture semantic meaning and relationships between words. To manage imbalance distributions, Yan *et al*. (2018) proposed a new model which combines Deep Boltzmann (Salakhutdinov and Hinton, 2009) with CNN using title and abstract with accuracy between 0.57 and 0.77 depending on the dataset. To raise accuracy up to 0.82, Burns *et al*. (2019) added evidence fragments (paragraphs which reference a figure), PubMed Medical Subject Headings, and captions, reaching 0.91 if including full text. Lee *et al*. (2018) used title, abstract, journal information, and publication type over three datasets with good precision (>0.91) and recall (>0.93) on the balanced datasets.

CNN and word embeddings are ideally suited to capture the spatial correlations and directional information required in specific types of literature mining, such as when searching for molecular interactions. Nonetheless, complex algorithms constitute a technological barrier, as they require computer programming and deep mathematics knowledge in addition to expensive hardware requirements. Moreover, they may not generalize well to diverse biocuration projects since they lose performance with disparate datasets. Thus, it remains to be determined if relatively simpler approaches than CNN and word embeddings may effectively solve the biocuration triage problem, especially in the common situation of strongly unbalanced datasets (Almeida *et al*., 2014).

NeuroMorpho.Org is a database of digitally reconstructed neuronal and glial morphology (Ascoli *et al*. 2007; Akram *et al*. 2022). The curators of this resource continuously search the peer reviewed literature and actively contact authors to acquire, process, and publicly release new data (Ascoli, 2015). The main bottleneck for the project is the triage task: recognizing relevant peer reviewed publications, which are 10% of the total set of retrieved documents, takes one skilled individual an average of 3.5 hours per article (Akram *et al*., 2018) in addition to the substantial time required for personnel training (Ascoli *et al*., 2017). We developed PaperBot (Maraver *et al*., 2019), a tool to improve the publication identification and acquisition process. Adoption of PaperBot drastically increased the number of found publications from an average of fewer than 800 per year to almost 6000, further aggravating the triage problem. A major challenge for NeuroMorpho.Org is that the key paragraphs used for triage are usually in Methods, Results or figure legends, making the use of full text necessary (Bijari *et al*., 2020). However, our experience with manual triaging led us to hypothesize that word order may be largely disregarded for this step; instead, word order becomes important for selecting the bio-entity data within the text once relevance is established. Based on these premises, here we propose a simple deep learning tool designed to be installed and used without machine learning knowledge. It trains quickly on any modern computer and potentially could be applied to projects with disparate relevance criteria where previously annotated text is available to re-train the classifier model. Specifically, the tool assesses whether the text is related to the domain of interest (neural morphology) or off-topic. If the text is related, the tool calculates the likelihood that the article describes digitally reconstructed neurons or glia. Moreover, the tool is easy to re-train for capturing new features or improving performance.

We have deployed and are actively using the tool to automatically discard non-relevant publications and sort the remaining ones by relevance. This helps us review first publications with higher likelihood of containing neural reconstructions, increasing the amount of data that become available to the research community.

## 2 Methods

We trained a deep learning classifier to return the likelihood that digital reconstructions of neuronal or glial morphology were performed during the research described in a text. The following subsections describe the steps followed to develop this smart tool (Fig. 1).

**Fig. 1.**
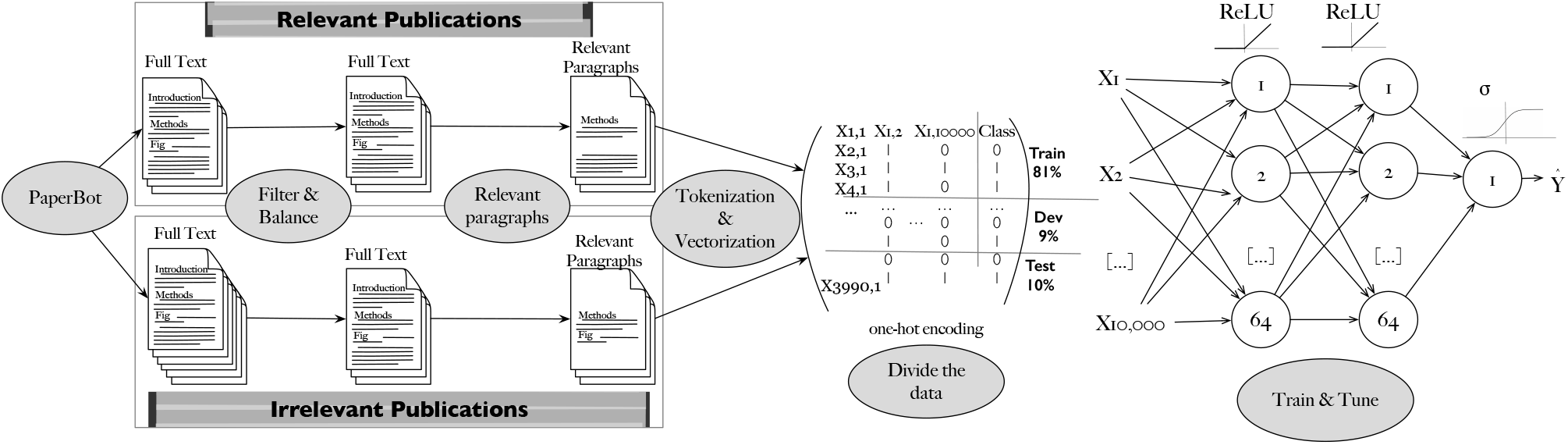
Schematic workflow in the operation of the deep neural network classifier. The first step is the data acquisition, performed using PaperBot, a tool to search peer-reviewed publications from multiple sources using keywords. These acquired publications are each manually annotated as relevant or irrelevant depending on whether they describe new neural reconstructions. The second step includes filtering away the publications extracted from pdf files (due to lower text quality), balancing relevant and irrelevant datasets, extracting relevant paragraphs from the text to reduce noise and improve performance, and tokenizing to vectorize the text. The final step divides the data into train, validation, and test sets to optimize the deep learning classifier hyper-parameters and evaluate performance.

### 2.1 Acquiring data: PaperBot

The pipeline starts by finding and acquiring all the new research publications that potentially describe digitally reconstructed neural morphology. For this task, we utilize PaperBot, a configurable, modular, open-source web crawler that automatically identifies peer-reviewed articles based on periodic full-text searches across publisher portals (Maraver *et al*., 2019). Queries rely on customizable combinations of keywords that, for this project, focus on the tracing systems (e.g., Neurolucida, Imaris, Simple Neurite Tracer, NeuronJ), file format extension (SWC), or terms related to neural reconstructions (skeleton, Sholl analysis, filament, morphology, dendrites, axons, confocal microscopy, etc.).

PaperBot acquires the full text of available publications including title, abstract, figure legends, and all sections such as Introduction, Methods, Results, Conclusion, References, and Supplementary Materials. From publishers that offer an Application Programming Interface (API), PaperBot gleans high-quality text made up of paragraphs. When only PDF files are provided, the extracted text is of lower quality as it contains page headers and footers mixed with the content as well as occasional split and incomplete words. We only utilized the API-derived high-quality text for training the classifier, but also included the PDF-derived lower-quality text for analyzing the results.

### 2.2 Filtering and balancing data

The second step consisted of selecting a pool of publications suitable for training the classifier based on existing annotations. Since inception of the NeuroMorpho.Org project, our manual curation process categorized every paper returned by keyword search with one of four labels: (i) relevant, if the text describes neural morphology that was digitally reconstructed in the course of the described research; (ii) linked, if the text describes neural morphology that had been digitally reconstructed previously as described in a cited reference; (iii) uncertain, if the text is insufficient for an expert reader to unambiguously ascertain whether or not neural morphology was digitally reconstructed; and (iv) irrelevant, if the text does not describe digital reconstructions of neural morphology. Given the goal of the classifier, we only included publications in the first and last category in the training dataset.

The resultant dataset consisted of 1,995 relevant and 2,971 irrelevant publications. We balanced this dataset by random under-sampling to avoid a bias towards the majority class as typically observed when learning from imbalanced distributions (Weiss and Provost, 2003). Random under-sampling discards stochastically selected entries from the majority class to match the sample size of the minority class, which works well in practice and is simple to implement (Johnson and Khoshgoftaar, 2019).

### 2.3 Selecting relevant paragraphs

Each publication contains many paragraphs with hundreds of words each. Only a minority of these paragraphs describe digital reconstructions of neural morphology. To reduce noise and improve performance, we removed from the input text all paragraphs that did not contain any of the keywords utilized in the original PaperBot query described in section 2.1 above. This process mimics the behavior of human curators performing triage, who search the text for keywords and then evaluate the corresponding paragraphs to ascertain the relevance of the publication.

### 2.4 Tokenizing and vectorizing data

Next, we convert the text of the selected paragraphs to numerical vectors for use in deep learning through the following steps using the spaCy library (Honnibal *et al*., 2020):

1. Split text into single words (tokenization).
2. Remove punctuation and stop words, which do not add context value.
3. Extract the lemma for each word, which is the meaningful root form of a term.
4. Convert all words to lowercase.
5. Out of the resulting vocabulary of ~ 170*k* words, selected the most frequent 10*k* to increases performance.

Vectorization then converts words into a numerical representation using one-hot encoding, which creates a vector containing 1 if the word is present or 0 if absent.

### 2.5 Dividing the data into train, validation, and test sets

We randomly split the data into two non-overlapping sets: one for train and validation, and the other for testing. The test set represents 10% of the data and is held out during training and validation to obtain an unbiased estimate of the performance. On the remaining 90% of the data, we applied 10-fold cross validation (Refaeilzadeh *et al*., 2009), where the dataset is split in two further subsets: one used to learn the model and the other used to validate it. During this step the data are split 10 times randomly and with each iteration a different set of data is held out for validation. The validation set, consisting of 9% of the original data, is used to tune the neural network and search the hyper-parameters yielding the best results for the model trained using the remaining 81% of the data.

### 2.6 Training the neural network model

The objective of a feedforward neural network *ŷ* = *f* (*x*; Θ) is to learn the parameters Θ that result in a good estimate function of the data *x* provided. The network architecture is formed by units grouped in layers connected in a chain structure, each layer represented by equation (1):

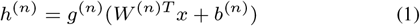

Here, *n* represents the layer which consist of many units acting in parallel, each representing a vector-to-scalar function; the transpose of matrix *W* provides the weights of a linear transformation along with the bias vector *b*; and the non-linear activation function *g* is applied to compute the values of the hidden layer. To learn a good estimate *ŷ*, the back-propagation step computes the gradient to minimize the deviation between the true value and the predicted one, called the loss function (*Loss*(*ŷ, y*)). For balanced-classification problems, accuracy (equation 2) is used to measure the performance of the classifier based on True Positive (TP), True Negative (TN), False Positive (FP), and False Negative (FN):

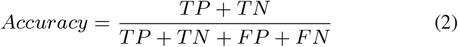

### 2.7 Selecting the hyper-parameters

Hyper-parameters are settings that control the behavior of the machine learning algorithm, but are not learned by the algorithm itself (Goodfellow *et al*., 2016). The goal is to minimize the loss function while maximizing accuracy.

There are two approaches to selecting hyper-parameters in a neural network model: manually when the computational resources are limited or automatically otherwise. We applied the manual approach: we fixed all values except one to be determined, starting from the most important parameters. In each case, we aimed to search through a broad enough range to find good results, but sufficiently contained to finish the task in a reasonable amount of time. To facilitate the process, we plotted accuracy and loss (Fig. 2) and selected the most promising value from the tested ones (Table 1). The order and role of the hyper-parameters is the following.

**Table 1.** Hyper-parameters used to train the deep neural network

| Hyperparameter | Applied Value |
| --- | --- |
| Hidden units | 64 |
| Hidden layers | 2 |
| Optimization algorithm | Stochastic Gradient Descent:<br>(1) $\hat{y} = \frac{1}{m} \nabla_{\theta} \sum_{i=1}^m L(x^i; \theta; y^i)$<br>(2) $\theta = \theta - \alpha \hat{y}$<br>Learning rate ($\alpha$): 0.01<br>Mini-batch size ($m$): 8<br>Loss Function ($L$): *Mean Square Error* $= (y - \hat{y})^2$ |
| Activation functions (g) | *Hidden layers* : $ReLU = \max(0, z)$<br>*Output layer* : $\sigma = \frac{1}{1+e^{-z}}$ |
| Epochs | Number of training iterations over the dataset: 30 |

**Fig. 2.**
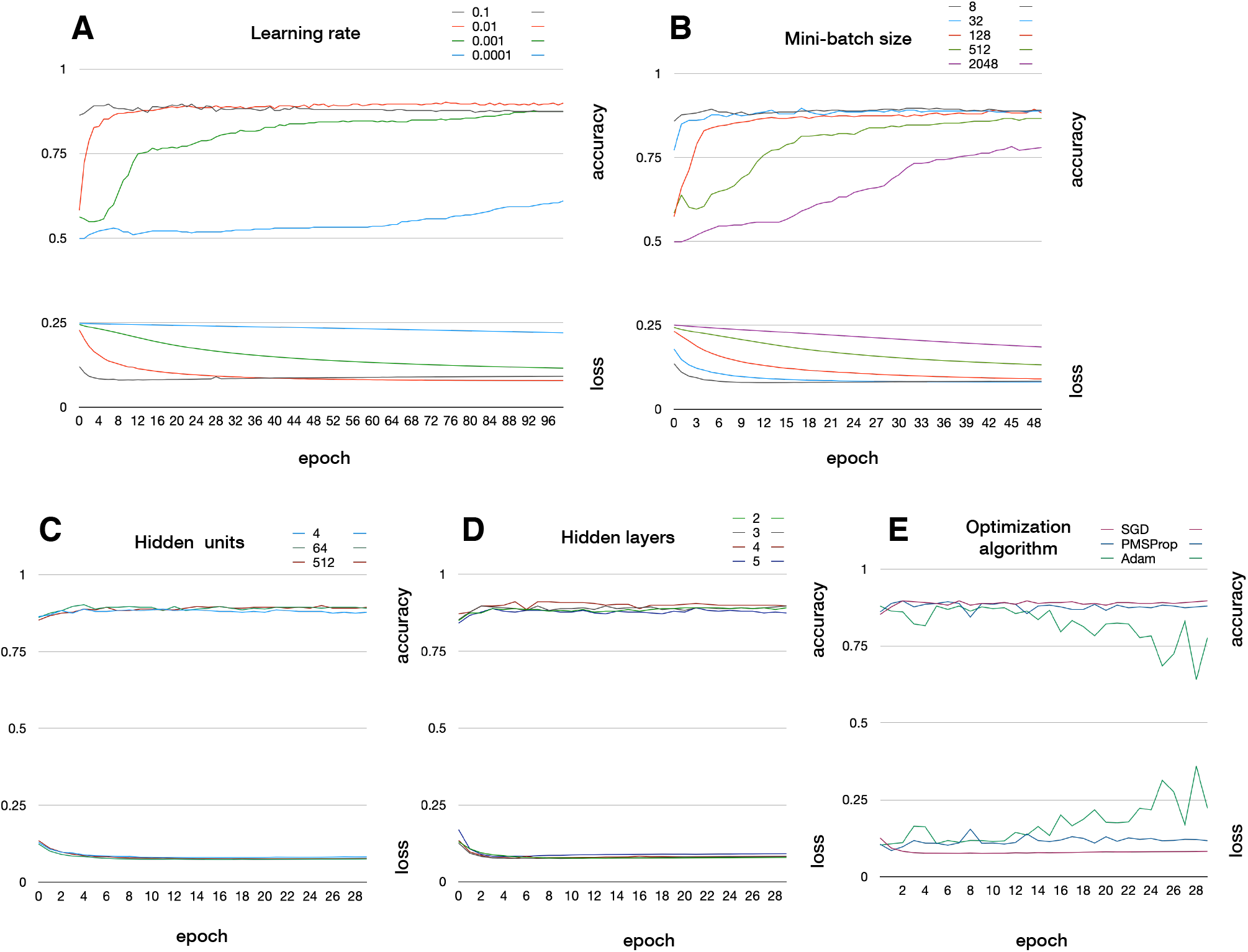
Hyper-parameters accuracy and loss drawings used to select the most promising values. (A) Learning rate. (B) Mini-batch size. (C) Number of hidden units per hidden layer. (D) Number of hidden layers. (E) Optimization algorithm: Stochastic gradient descent (SGD) (Ketkar and Ketkar, 2017); adaptive gradients methods RMSProp (Tieleman and Hinton, 2012), and Adam (Kingma and Ba, 2014).

1. The learning rate is the fractional amount by which the weights are updated during training and controls the training speed; a low rate typically yields results closer to optimum at the cost of a slow process, while high rates can produce faster learning, with the risk of non-monotonous (e.g., oscillatory) convergence.
2. The model weights are updated by gradient descent after each batch of data. Using the entire training set as batch requires high memory, while using a single stochastically sampled element is noisy and inefficient. The mini-batch gradient descent size is a value between 1 and the whole training dataset.
3. The number of hidden units form the width of the neural network. Increasing this number typically improves the model representation while increasing computational cost.
4. The number of hidden layers represent the depth of the neural network. Increasing this numbers also typically improves the model representation while increasing computational cost.
5. The optimization algorithm finds the optimal values for the weight vectors.
6. The recommended activation function for modern neural networks is the rectified linear unit (ReLU) (Jarrett *et al*. 2009; Nair and Hinton 2010; Glorot *et al*. 2011), defined as the positive part of the argument, because it generalizes robustly and works effectively with gradient descent algorithms (Goodfellow *et al*., 2016)
7. The epochs are the number of iterations performed over the entire training set. We observed that our neural network model reaches stable results after 30 epochs.
8. The loss function measures how close the estimated output is compared to the input class value.

### 2.8 Selecting the thresholds

The decision or classification thresholds are the likelihood values selected to decide the belonging to a given class. We defined three thresholds identifying four different categories: *Positive High* publications are directly accepted as relevant; *Positive Low* and *Negative Low* publications are reviewed by curators to check the relevance classifier label; and *Negative High* publications are automatically discarded.

### 2.9 Coding the tool

We developed the code in the Python programming language. To create the model, we used the deep learning open-source framework Keras (Chollet *et al*., 2015), which runs on top of TensorFlow and facilitates its use. TensorFlow is a deep learning library developed and maintained by Google that provides low and high-level APIs built to run on multiple CPUs or GPUs (Li *et al*., 2019).

The tool is containerized with Docker as a service and exposes a REST API that offers three services: Train, Classify, and Search Keywords.

The Train service reads all files storing the training text instances, vectorizes the texts, and trains the classifier. Two new files are created as a result of the classification: the tokenizer vocabulary and the classification model. Both files will be used in the classification step to vectorize new data and predict the relevance. The output returns data in json format containing the accuracy and loss values for the train and test sets, the epochs or training iterations, the number of train and test samples, the version of the model used when classifying (in case multiple versions are maintained and evaluated), and the classification result for each of the test samples [true value, classification value] (Fig. 3A).

**Fig. 3.**
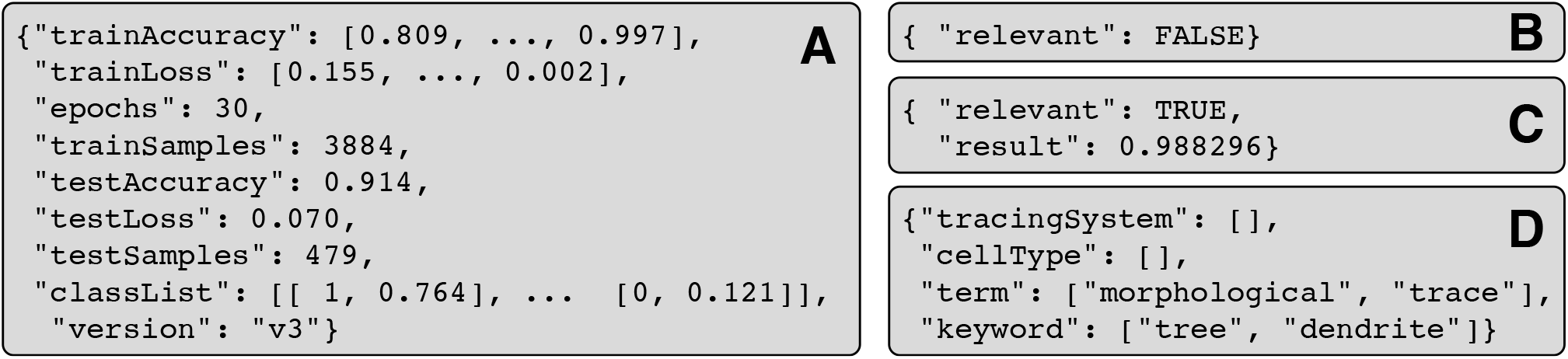
API json results for each of the calls. (A) Train service. (B) Classify service - example of irrelevant publication. (C) Classify service - example of relevant publication, with likelihood. (D) Search Keywords service.

The Classify service returns json format data with the likelihood of relevance of the given text. First, it searches for the relevant paragraphs on the text; a paragraph is relevant if it contains any pre-defined keywords. Occasionally Google Scholar returns publications that do not contain any of the query keywords, an unintended side effect of its expanded semantic search approach; our tool identifies these cases as off-topic and classifies them as irrelevant (Fig. 3B). If there are relevant paragraphs, their text is vectorized using the tokenizer file created in the previous step and sent to the classifier to predict the relevancy likelihood (Fig. 3C) using the model file also created in the previous step.

The Search Keywords service selects all instances found in the text from the user-configurable set of keywords by extracting the token and comparing the lemma. The result is a json data format object composed by each of the file names and the list of the found keywords (Fig. 3D).

## 3 Results

PaperBot was installed in January 2016 and tested during a two-year period. It was then launched retroactively to search automatically what was previously queried by hand, and found several missed publications from previous years. In 2018 the set of keywords used to find publications was vastly increased taking advantage of the automated processing capability. Once launched, it outperformed the manual task in the current years while still finding missed publications for previous periods (Fig. 4a). On the downside, the number of irrelevant publications found increased accordingly (Fig. 4b).

**Fig. 4.**
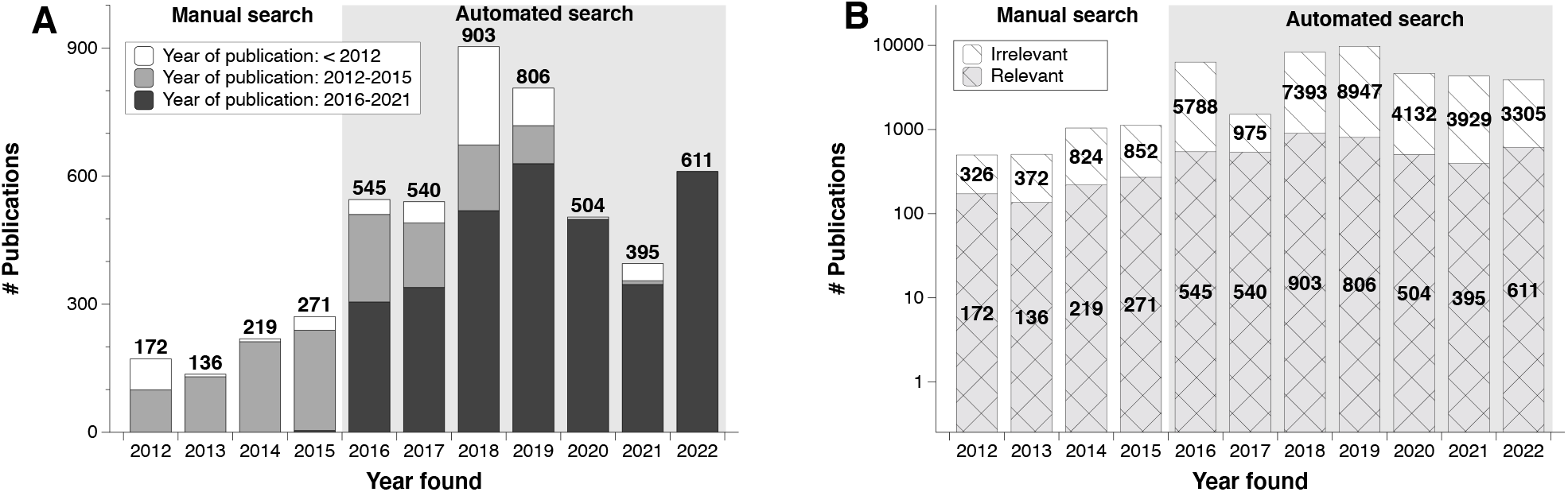
Number of publications found using manual (clear background) and automated (gray background) searches. (A) Relevant publications by the year in which they were found grouped by published period: before 2012 (white stacked bars), 2012-2015 (light grey), and 2016-2022 (dark grey). (B) All publications by the year in which they were found grouped by relevancy: relevant (double cross pattern) and irrelevant (single line pattern).

We trained and tuned the deep neural network described in the Methods (Fig. 1) until its performance fulfilled our lab requirements: an average accuracy greater than 93% minimizing loss (Table 2). Correctly and incorrectly labeled results for the test set are shown as a confusion matrix in Table 3.

**Table 2.**
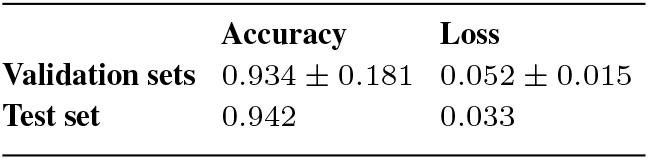
Performance of validation and test sets

|  | Accuracy | Loss |
| --- | --- | --- |
| Validation sets | 0.934 ± 0.181 | 0.052 ± 0.015 |
| Test set | 0.942 | 0.033 |

**Table 3.**
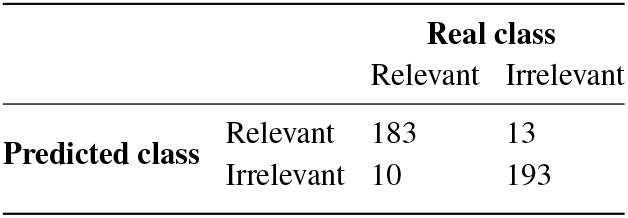
Confusion matrix of the test set

Approximately 10% of all the publications found are relevant (between 9% *−* 13% depending on the year: Fig. 4b). Triage is an intrinsically probabilistic operation, as each publication is assigned a relevancy likelihood. Thus, choosing different thresholds for NeuroMorpho.Org depending on the damage of mislabeling proved to be a successful strategy. Mistakenly labeling a relevant article as irrelevant is harmful for the project, because it means the reconstructions cannot be requested and will be effectively lost. Conversely, incorrectly labeling an irrelevant article as relevant only wastes labor since we request the data and then correct the records manually when the authors reply indicating that no relevant data were collected. Therefore, instead of using a traditional binary classification (relevant vs irrelevant), we convert the probability returned by the classifier into four labels, adding low confidence and high confidence to each of the classes as described in section 2.8. To select the most appropriate values for each of the thresholds we calculated the percentage of labor saved from curators against the percentage of incorrectly classified publications (Fig. 5). Specifically, we selected the 0.18 threshold as Irrelevant with high confidence, which saves 96.4% of manual processing time with only 3.10% of misclassified articles. At the opposite end, we selected a threshold of 0.98 as Relevant with high confidence, which saves 83.06% of manual processing time while misclassifying just 0.97% of articles. After these fully automated steps, 3.63% of low-confidence irrelevant articles (probability between 0.18 and 0.5) and 16.94% of low-confidence relevant ones (probability between 0.5 and 0.98) remain for manual review. On average, with the selected thresholds, from a set of 100 publications identified by PaperBot, only 5 need to be reviewed, of which 2 will be relevant and 3 irrelevant. From the remaining 95 not reviewed, 3 publications will be miss-classified (Fig. 4).

**Fig. 5.**
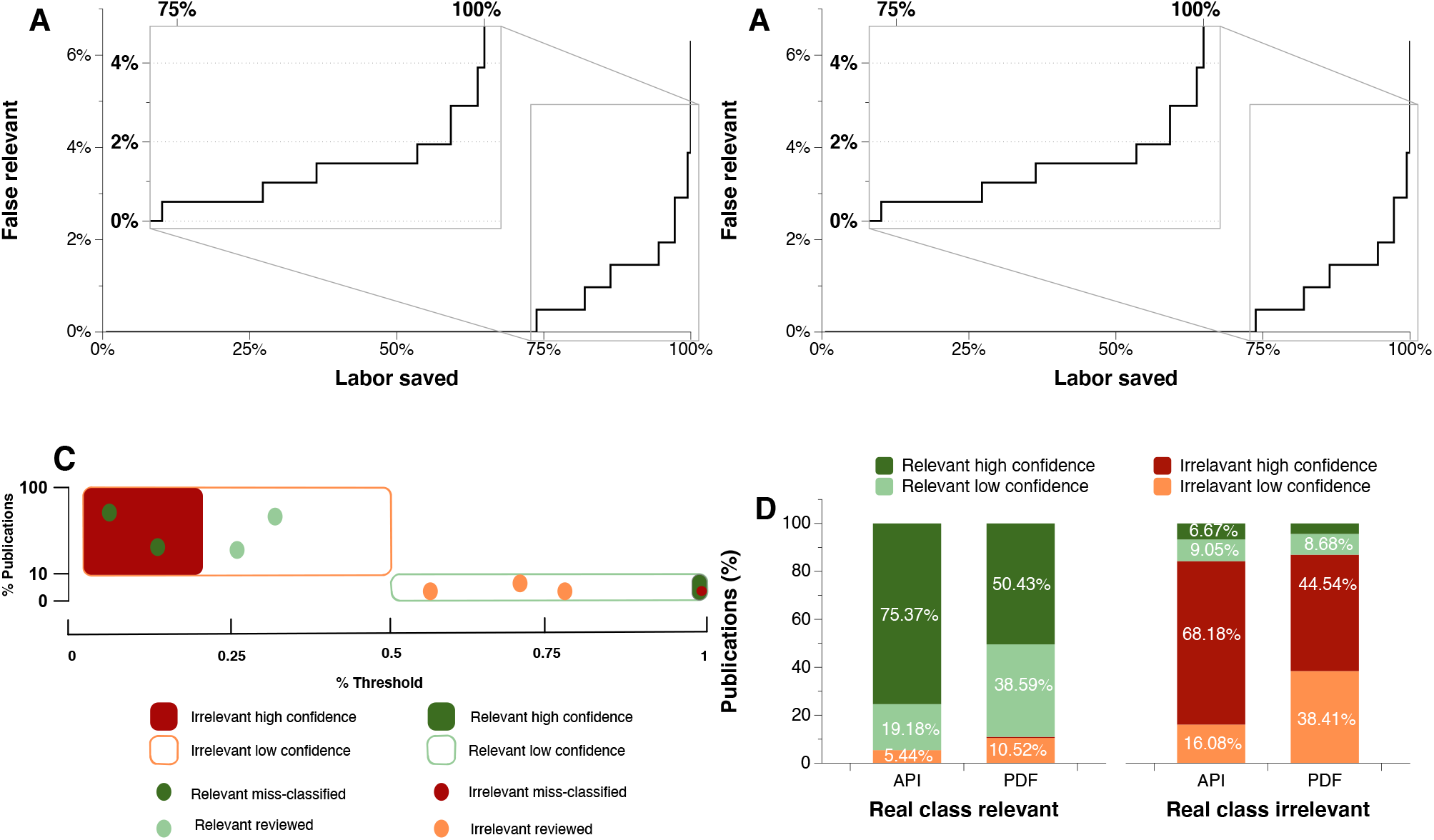
Selecting the most convenient thresholds. (A) Proportion of irrelevant articles incorrectly classified as relevant (false relevant) as a function of the fraction of labor saved by accepting the automated classification without review. The inset displays the enlarged range between 75% and 100% of saved labor. (B) Proportion of relevant articles incorrectly classified as irrelevant (false irrelevant) as a function of the fraction of labor saved by accepting the automated classification without review. The inset displays the enlarged range between 75% and 100% of saved labor. (C) Using test labeled data we select optimal thresholds to maximize saved labor while minimizing misclassification errors and the number of publications to be manually reviewed. (D) Once the classifier is deployed, we analyze the results by type of text: high-quality text obtained from publishers’ APIs and low-quality raw text extracted from PDFs.

The final version of the classifier has been operating in the lab since November 2020 in review mode, where all the publications are reviewed by an expert after being labeled by the classifier. Most of the publications are correctly labeled by the classifier thanks to the low loss obtained for the classification model.

In the real environment, PaperBot retrieves both high-quality text from the publishers’ API, which represents 62% of the total, and lower-quality content extracted from pdf files. This remaining 38% was not used for model training but is still included in the analysis. As expected, when separately analyzing the model on high- and low-quality text, the latter results in lower performance. The Relevant with high confidence publications represent 75.37% of the high-quality text pool, but only 50.42% for the low-quality text pool, implying a 25% increase in articles requiring review. Similarly, the Irrelevant with high confidence publications are 68.18% of the high-quality texts, but only 44.54% of the low-quality ones, increasing the need for manual review by 23.64% (Fig. 4).

From the practical viewpoint of NeuroMorpho.Org, the most important success metric for the relevancy classification is the number of mined reconstructions and corresponding availability of shared data. From this perspective, we can compare the number of reconstructions mined and shared by the authors during three main time periods: manual search and evaluation, automated search and manual evaluation, and automated search and evaluation (Fig. 6). Although during the automated search many publications were found, it has been impossible to process all of them because of the human resources required to manually evaluate the publications.

**Fig. 6.**
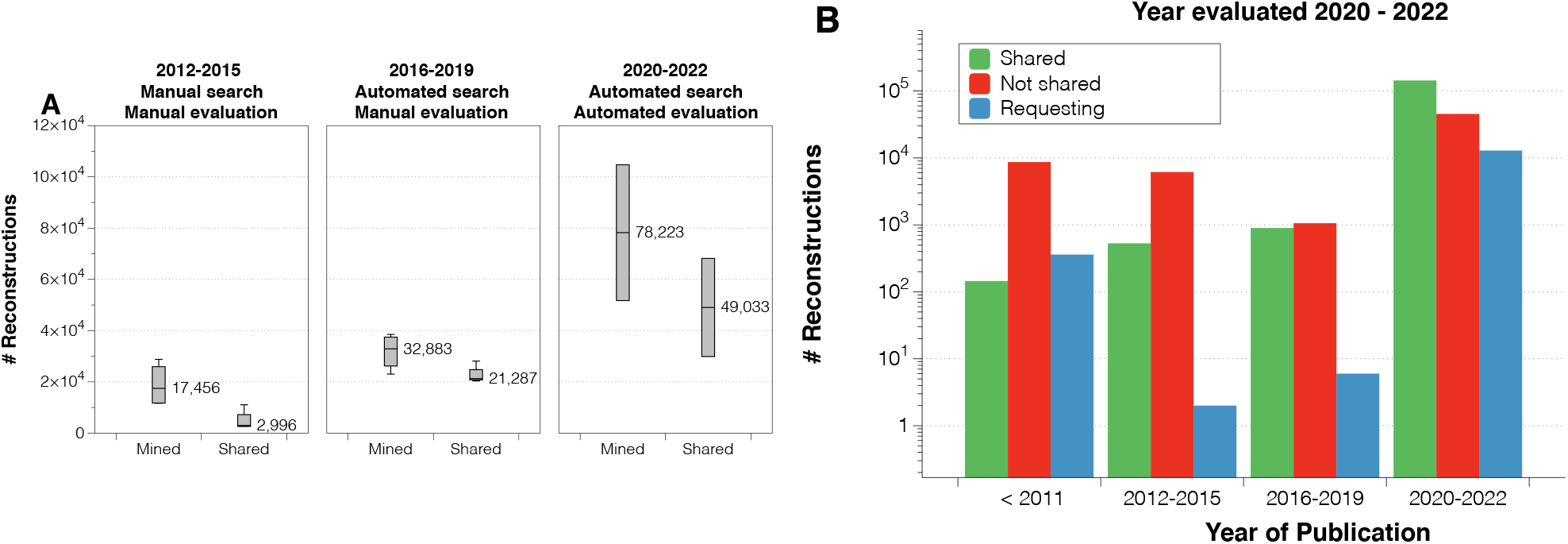
Numbers of mined and shared neural reconstructions. (A) Number of mined and shared reconstructions by year of evaluation for three periods: manual search and evaluation (2012-2015), automated search and manual evaluation (2016-2019), and automated search and evaluation (2020-2022). In addition to the more than tripled mined volume between the first and third period, note that only ~35% of mined data were shared from the first period and more than 80% in the last period. (B) Number of reconstructions shared (green), not shared (red), and pending or being requested (blue) to determine the availability for the evaluation years 2020-2022 grouped by published date.

Once the automated evaluation was deployed, we were able to process all of these publications, increasing by 2.37-fold the yearly average reconstructions mined compared to the previous period (Fig. 6A).

Interestingly, we found that requesting data on time improves the shared rate (Fig. 6B). We are currently requesting data for old publications that were missed during the manual search period. However, this attempt is often frustrated by many challenges: the contact information may be outdated, essential lab personnel may no longer be available, or the data may be lost due to storage obsolescence.

From a technological standpoint, using a simple neural network approach has allowed us to train, test, and tune many parameters very efficiently. Reading the data from pre-processed files and performing 30 training iterations for 10 k-fold takes 2.5 minutes on a modern GPU laptop. Most important, the classifier evaluates one publication every four seconds. One person manually searching publications online, finding the full text, reading it, and assessing the relevance spends 3.5 hours per relevant publication found. During the testing period of the tool described here, this time went down to 1.68 hours, which we expect to further decrease when we stop reviewing and confirming the classifier results.

## 4 Conclusions

The number of peer-reviewed publications continues to grow every year, making biocuration an arduous task; for scientists in general to keep up to date on new research, the step of triage must be automated. For the specific use-case of NeuroMorpho.Org, ordering all articles by relevance likelihood helped prioritize the data requests starting from the publications that were most likely to contain reconstructions, which increased the sharing rate. Timely identification is essential because delayed data requests were shown to be less frequently fulfilled.

Triage remains an open problem, because different labs typically develop ad-hoc solutions tailored to their own personalized needs. Thus, the resultant tools tend to only perform well for the original datasets for which they were designed. Conversely, the services we introduced here could be scalable to different biocuration projects with alternative definitions of relevancy. While we demonstrated the successful application to NeuroMorpho.Org, the same strategy could be expanded in many other triage applications (Anderson *et al*., 2021).

Deep learning has released curators in our lab from non-creative work resulting in a better use of our time and saving almost 2 hours per relevant paper reviewed. While we have so far selected very conservative thresholds, relaxing them could save even more resources at the cost of marginal data loss.

A potential direction to extend this line of research is to combine and integrate the described work with the recently introduced automated extraction, from the same articles, of rich metadata pertaining to the identified reconstructions, such as the animal species, sex, and age; the brain region and cell type; and the histological and imaging protocols (Bijari *et al*., 2022). At the same time, it may be worth exploring in future research the possibility of combining full text with figures (Jiang *et al*. 2022; Ljungquist *et al*. 2022), morphological metadata, and bibliographic information (such as journal and authors), to learn new models and improve the results further.

## Acknowledgements

We acknowledge all the curators that had labeled huge amounts of data, which was key for the success of the project. Special thanks to Masood Akram who reviewed and confirmed the accuracy of the evaluations, and to Kayvan Bijari for feedback and advice on natural language processing and deep learning.

